# homeRNA Just Dropped! Streamlining Remote Blood Collection and RNA Stabilization for Smaller Sample Volumes with homeRNAdrop

**DOI:** 10.64898/2026.09.22.753625

**Authors:** Madeleine Eakman, Filip Stefanovic, Winston Liang, Ingrid H. Robertson, Aatman Jindal, Karen N. Adams, Erwin Berthier, Amanda J. Haack, Ashleigh B. Theberge

## Abstract

At-home sampling can address obstacles to participation in transcriptomics research. We previously developed the homeRNA and homeRNAmax platforms; these custom-designed tubes containing RNA stabilizer interface with a blood tube compatible with upper-arm blood collection devices, allowing for blood collection and RNA stabilization to occur from a participant’s home. In this work we introduce homeRNAdrop, an insert for a commercially available blood tube (BD Microtainer) that divides the tube into two compartments: the top compartment holds 200 μL blood and the bottom compartment holds ∼590 μL liquid stabilizer (RNAlater). By combining blood collection and RNA stabilization into the same tube, homeRNAdrop streamlines sample preparation for participants and allows for more compact sample storage for the lab. In a remote pilot study, n=25 participants from across the United States tested the homeRNAdrop kit. The n=23 samples successfully received by the lab show that samples collected using homeRNAdrop are of sufficient RNA quality (all RINs > 6, a common cutoff for analysis) and yield (mean yield=1.44 μg) for downstream gene expression analysis. Almost all participants who successfully collected blood using the Tasso Mini device found the homeRNAdrop device easy to use (n=23/24). Overall, this work adds a simplified, user-friendly platform to the growing suite of tools for remote blood RNA stabilization.

## Introduction

Remote studies are rapidly emerging as the new standard in clinical and transcriptomics research, in part because they eliminate the need for participants to be physically present at a study site^1^. In particular, blood sample collection can be challenging, because venous blood draws require a highly trained phlebotomist and can limit the opportunities for sample collection^2^. Dried blood spot (DBS) sampling is an existing remote sampling technology^3–12^ that can be used for gene expression analysis^13,14^, but a stabilized liquid blood sample with a larger volume of blood (>100 μL) could allow for more types of downstream analyses to be performed. To this end, we developed custom-designed tools that leverage the Tasso upper arm blood collection device to allow for immediate stabilization of blood RNA in remote studies^15,16^. Called homeRNA, these platforms have been successfully used in studies investigating gene expression changes related to SARS-CoV-2 infections^17,18^ and wildfire smoke exposure^19^, have captured an induced immune response comparable to PAXgene-stabilized venous blood–the current gold standard in transcriptomics research^20^, and has been shown to be usable in many warm climates through real-world shipping tests^21^ and in-lab experiments^22^, as well facilitating increased engagement in longitudinal studies^23^.

While homeRNA platforms are robust tools for stabilizing RNA in home-collected blood samples, laboratories running studies that do not require a full mL of blood might appreciate a smaller device that does not require a separate tube for analyte stabilization. In this work we present a new embodiment, homeRNAdrop, an insert for the BD Microtainer which allows for blood collection and RNA stabilization to occur in the same tube, simplifying sample collection for the user and kit assembly for the study team, and making long-term sample storage more efficient compared to our previous homeRNA devices by reducing the footprint in a -80 °C freezer.

## Experimental Details

### Design process for insert

The homeRNAdrop insert is designed to separate the BD Microtainer tube into two compartments, where the bottom compartment holds up to 590 μL of liquid stabilizer and the top holds up to 200 μL of blood, allowing users to collect the blood sample and stabilize the RNA within the same tube. Design iterations were developed on Solidworks and 3D printed using a Form 3B 3D printer (Formlabs). The insert design also features a cone channel separating the two compartments (Figure 1B) to prevent contact between the user and the RNAlater during stabilization. These designs were tested for leakage and spillage using a simulation of the mechanical agitation of the shipping process similar to that described in Eakman, Stefanovic et al. (2026)^16^; five BD Microtainers with the homeRNAdrop inserts in place were filled in the bottom compartment with 590 μL of RNAlater, then they were placed in the shipping box and dropped from 1.2 m under the following conditions (one sample per condition): 1) flat on the bottom, 2) flat on the top, 3) flat on the long side, 4) flat on the short side, and 5) on a corner. Drop test procedures are in accordance with 49 CFR 178 subpart M.

**Figure 1.**
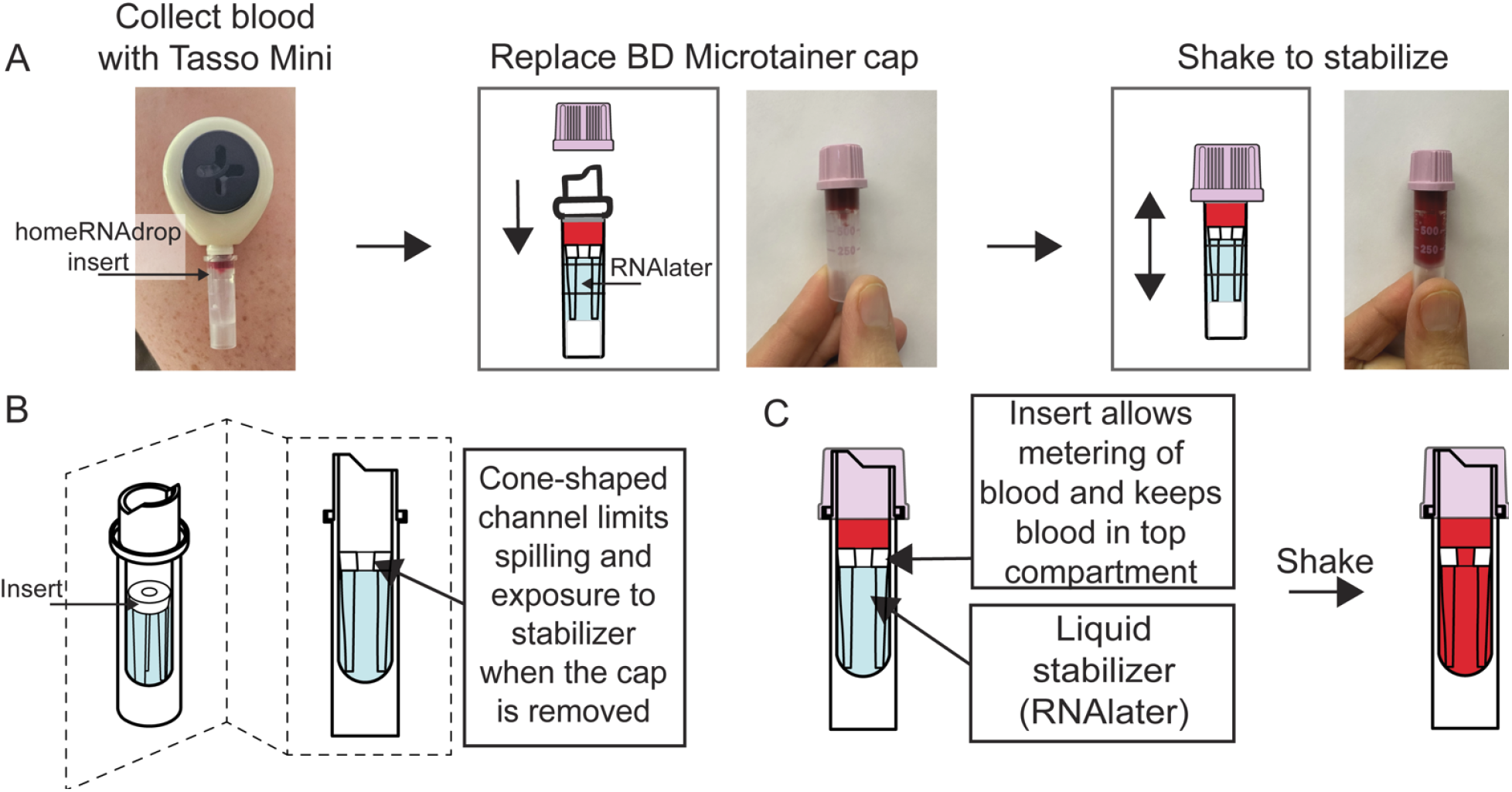
Workflow and features of homeRNAdrop. A) Workflow for using homeRNAdrop. Users collect blood using the directions for the Tasso Mini blood collection device, replace the blood tube cap, then shake the blood and stabilizer fluid together to stabilize. B) Schematic of BD Microtainer with homeRNAdrop insert and stabilizer fluid. C) Cross-section schematic of blood-filled homeRNAdrop before and after shaking to stabilize.

Designs that passed these in-lab tests were then subject to a real-world shipping test to ensure minimal leakage during the shipping process; homeRNAdrop devices were assembled, filled, weighed, and placed in a homeRNAdrop kit. These kits were shipped to a lab member’s house in the Seattle area, and shipped back to the lab. The homeRNAdrop devices were then weighed upon receipt. Fluid that had leaked to the top compartment was then removed with a pipette before weighing again; the mass difference was then used to calculate the fluid that leaked using the measured density for RNAlater 1.2125 g/mL. Shipping tests were performed in duplicate, where two replicates were sent in separate packages to one lab member’s house through UPS and another two were sent to a different lab member’s house on the same day. Later iterations of this test also included bubble wrapping the cardboard Tasso box; while the usefulness of bubble wrapping is inconclusive from our shipping tests, we continued to use bubble wrap in the pilot study as a further safeguard against impacts in shipping (see Fig. S1 and Table S1). Designs were considered to have passed the shipping test if the average fluid loss into the top compartment of the BD Microtainer among the packages shipped to both locations (total of four packages bubble wrapped and four packages non-bubble wrapped) was < ∼80 μL, and in no case did stabilizer fluid enter the cap. We note that since no stabilizer fluid entered the cap, any stabilizer fluid that enters the top compartment would still be available for stabilizing blood, but would also decrease the volume of blood the top compartment of the BD Microtainer can hold. Importantly, in this study we found that 20/23 samples had RNA yields above 500 ng and 3/23 samples had RNA yields above 100 ng, which are both sufficient for downstream analysis. (Engineering drawings for the homeRNAdrop insert design used in this work can be found in Fig. S2).

### Fabrication of the homeRNAdrop device

homeRNAdrop inserts are 3D printed on a Form 3B 3D printer (Formlabs) using clear resin. Inserts are then cleaned with the Formlabs Formwash in isopropyl alcohol (IPA) for 20 minutes, followed by a second wash in a separate Formwash filled with IPA for 10 minutes to ensure the inserts have no uncured resin. Inserts are then cured in a Formcure for 30 minutes at 60 °C in accordance with the manufacturer’s directions. Inserts are then placed in BD Microtainer tubes, and filled in the bottom compartment with 590 μL of RNAlater (Thermofisher) as the stabilizing reagent, and capped.

### Evaporative loss from homeRNAdrop

Ten homeRNAdrop devices were assembled according to the section above (see *Fabrication of the homeRNAdrop device*) before being filled with 590 μL of RNAlater. The devices were then capped and their initial mass was recorded. The filled homeRNAdrop devices were then placed in a microcentrifuge tube rack and left at ambient room temperature (∼21-24 °C). The mass of the homeRNAdrop was then recorded once per week for a total of 12 weeks. The change in mass from initial mass is calculated and converted to volume, using a density of 1.2125 g/mL for RNAlater. The resulting volume is then subtracted from 590 μL and converted to a percentage, representing the percent RNAlater loss from evaporation for each tube at each timepoint (Fig. S3).

### User experience pilot study

This study is IRB-approved under STUDY00007868 under the University of Washington IRB. Study procedures are adapted from Eakman, Stefanovic et al. (2026)^16^: for the usability pilot study, we recruited participants using Facebook and Instagram ads. We selected individuals who had not previously seen or used a Tasso device (by self-report) and who are not affiliated in any way with the Bioanalytical Chemistry for Medicine and the Environment (BCME) lab (Theberge lab). Individuals were eligible if they were healthy, adults aged 18 or older, and able to have lancet blood sampling (collected by self-report). People who met these criteria were then screened for age, gender, race and ethnicity, and state of residence. Balancing for these factors, 30 people were selected and sent an email with a link to the informed consent form (ICF). Upon completion of the ICF, participants would be considered as “enrolled” and were instructed to complete a survey with their shipping information. The study kit consisted of (1) instructions for sampling and sample return (Fig. S4-S5), (2) a Tasso Mini, (3) a Tasso warming pad, (4) an alcohol swab, (5) a bandaid, and (6) a homeRNAdrop device filled with 590 μL of RNAlater on the day of shipment (image and table of kit components can be found in Fig. S6-S7 and Table S2). The kits were wrapped in bubble wrap as this was found in pilot testing to further prevent stabilizer leakage into the top compartment, and packed inside a UPS LabPak alongside another, pre-labeled UPS LabPak for sample return. The packages were sent to the 27 participants who completed the ICF and shipping information form using the UPS next day air service and the tracking information was directly shared with the participants. Once a package was delivered to a study participant, another email with study instructions (see SI) was sent that included a link to an instructional video prepared by our lab. The participants then completed their sampling according to the instructions and completed a short usability survey (see SI); 25 participants completed this step. The sample was then sealed back in the Tasso box, placed into the return shippingLabPak, and kept indoors overnight. The following morning, participants left their samples at the designated pickup location and the study team scheduled a UPS pickup. Once the samples arrived back to the lab, they were stored at -20 °C until further processing.

In the pilot study outlined above, participants self-collected blood using the commercially available Tasso Mini device. The Tasso Mini device uses the BD Microtainer for blood collection, which comes in a variety of coatings. In the present manuscript and study, we opted to use the dipotassium ethylenediaminetetraacetic acid (K_2_EDTA) coated BD Microtainer tube to limit blood coagulation prior to stabilization; homeRNAdrop inserts were placed into the BD Microtainer, and the bottom compartments are filled with RNAlater prior to shipment to participants. Participants were instructed to collect blood for five minutes or when the blood reached the collar of the BD Microtainer (200 μL), whichever happened first; see instructions for use (IFU) in the Supplemental Information (Figure S3) for more information. Once the blood was collected, participants recap the BD Microtainer and shake it to mix the blood and stabilizer. These samples were then packaged and returned to the central lab as described above.

### RNA isolation and analysis

Sample RNA extraction was performed using the blood RiboPure RNA Purification Kit (Invitrogen) and following the manufacturer suggested protocol, with the exception that the two elutions in the final step were collected in separate tubes to increase the RNA concentration of our samples. The resulting RNA was then analyzed using the Agilent 2100 Bioanalyzer (Agilent Technologies, Inc.) to verify RNA integrity, and the RNA yield was measured on the Qubit Flex Fluorometer (Thermofisher Scientific).

## Results/Discussion

Previous homeRNA platforms^15,16^ consisted of a custom-designed stabilizer tube that houses a liquid stabilizing reagent (we use RNAlater), which users connect to a blood tube and shake to combine the blood and stabilizer, stabilizing the sample. This sample can then be shipped back to the lab for analysis. By contrast, homeRNAdrop allows for blood collection and RNA stabilization to occur in the same tube; participants only need to collect blood using a Tasso Mini upper arm blood collection device using a BD Microtainer with a homeRNAdrop insert inside, replace the blood tube cap, and shake to combine the blood and stabilizer (Fig. 1A).

The insert for the homeRNAdrop device is designed to divide the BD Microtainer blood tube into two compartments separated by a cone-shaped channel, which prevents the stabilizer from spilling out of the tube (e.g., when shipping, if the tube is accidentally knocked over by a participant) (Fig. 1B). The top compartment holds 200 μL blood, and the bottom compartment holds 590 μL liquid stabilizer (Fig. 1C), as dictated by the height of the insert. This ratio was selected to maximize the amount of blood that can be collected and stabilized using homeRNAdrop, while also ensuring adequate sample stabilization by including an additional 70 μL stabilizer volume compared to the manufacturer’s recommended ratio of 1:2.6 blood:RNAlater, accounting for any additional drops of blood that might be collected. However, this ratio could easily be changed (e.g., to accommodate a different stabilizer) by adjusting the height of the insert legs.

The dimensions of the cone-shaped channel were designed to minimize spilling upon impact while still allowing for mixing to occur between the two compartments. The mechanism of the spill-resistant feature has been described in our previous work^16^. The diameter of the cone was also designed to allow for a 14 gauge plastic needle to reach the bottom of the tube; using a plastic needle as opposed to the plastic Pasteur pipettes used in our other homeRNA platforms^15,16^ allows the channel to have a smaller diameter to further limit spilling, while avoiding the risks associated with metal needles.

The BD Microtainer was chosen as the blood tube for homeRNAdrop for broad compatibility with current upper arm blood collection devices; we use the Tasso Mini for the study in this manuscript, but any blood collection device that uses the BD Microtainer as its blood tube could be used, including the Tasso+, TAP Micro Select® (YourBio Health), and RedDrop One (RedDrop Dx). We note that it is also possible to refit this design to fit other tubes that are not the BD Microtainer.

To validate the efficacy of the homeRNAdrop, we conducted a pilot study recruiting participants from across the United States. Out of the 27 participants who completed the informed consent form (i.e., enrolled) and submitted their shipping information, 25 completed the study procedures and are reported in this work; these participants were recruited from 24 states (Fig. 2A), and the demographics roughly matched the US census (Fig. 2B), with an approximately even sex and age distribution (Fig. 2C).

**Figure 2.**
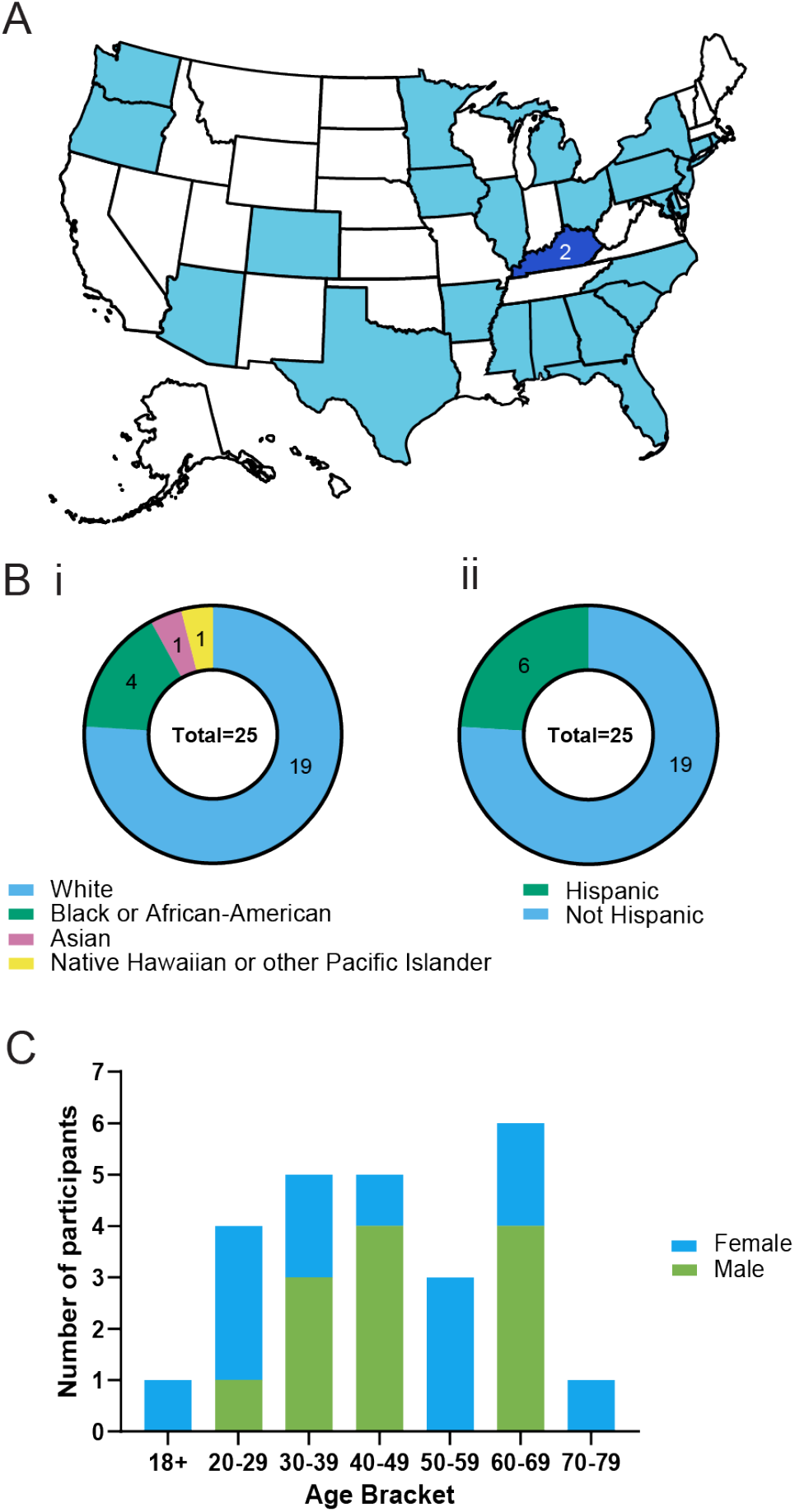
Summary of pilot study participant demographics. (A) United States map showing distribution of participants across 24 states. Each shaded state represents one participant except Kentucky, where two participants were recruited. (B) Distribution of races and ethnicities of participants per NIH categories. (C) Participant sex and age bracket.

Following recruitment, participants were sent homeRNAdrop kits. Each participant was asked to collect and stabilize one sample to capture the experience of a first-time user. The kits included detailed instructions for use (see SI) with an instructional video. Immediately following the usage of the kit, electronic surveys about the sampling and user experience were administered via REDCap. Participants typically reported no pain (n=20/25), with some (n=5/25) reporting mild pain (Fig. 3A). The participants also found the Tasso Mini device very easy (n=19/25) or somewhat easy (n=6/25) to use, with all but one able to collect blood. Of the participants who were able to successfully collect blood, most found the mixing process either very easy (n=20/24) or somewhat easy (n=3/24), with only one participant reporting the process to be somewhat difficult. Upon examination of the survey response, the participant who reported issues with the mixing stated that they had trouble understanding how to re-cap the BD Microtainer and that mixing took slightly longer than they had expected (the IFU states mixing generally takes about 10 seconds); these can be clarified for future studies with minor edits to the instructions for use. Finally, the majority of participants only needed up to 10 minutes to use the kit (n=19/24). Overall, the participant responses from the pilot study show that homeRNAdrop sampling is quick, easy to use, and painless, making homeRNAdrop a viable tool for application in remote clinical research.

**Figure 3.**
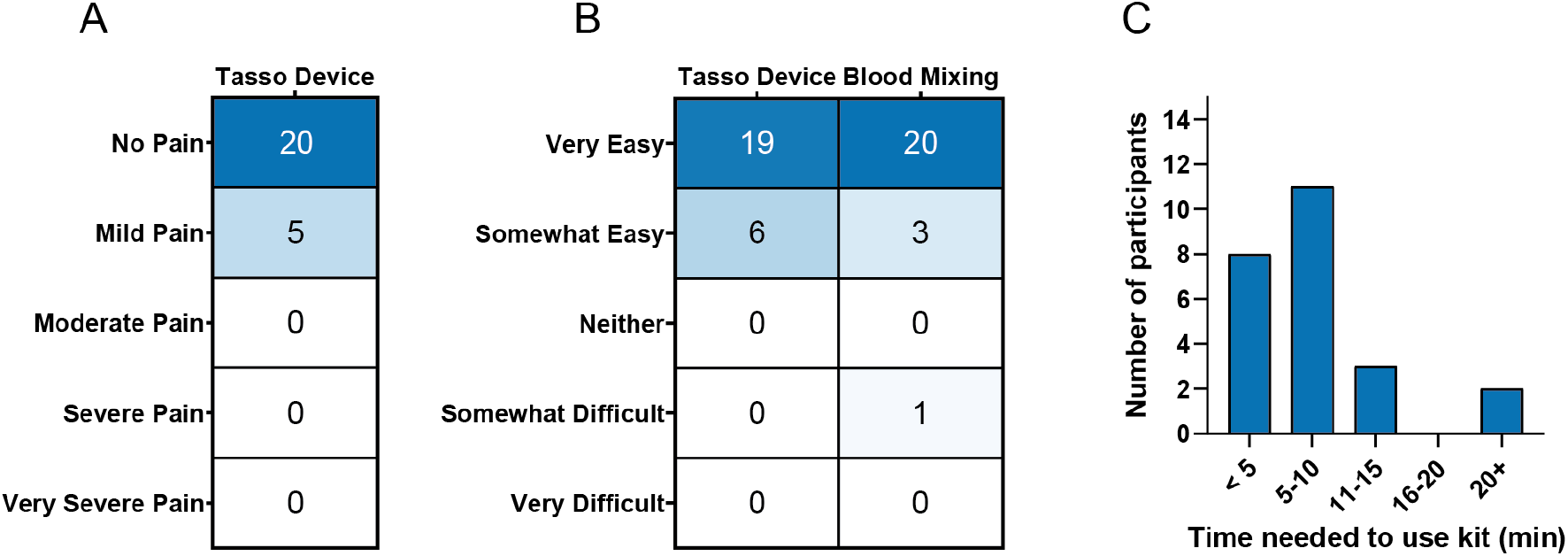
Participant-reported homeRNAdrop pilot study usability data. (A) Pain level by each participant following the use of the Tasso Mini device. (B) Participant-reported usability of the Tasso Mini and blood stabilization step of the homeRNAdrop device. Most (n=24/25) participants were able to collect blood and rated ease of blood mixing; one participant was not reported because they were unable to collect blood using the Tasso Mini device. (C) Time needed to use the kit, including reading instructions, collecting and stabilizing the sample, and packaging the sample for return shipping back to the lab. The participant who was unable to collect blood was not asked to report total time to use the kit.

Following the return of samples to the lab, RNA was extracted from the homeRNAdrop-stabilized blood samples with the RiboPure RNA Purification Kit using standard protocol. The extracted RNA was then analyzed for integrity and yield, as established in previous publications^15,16^.

Integrity analysis was performed using an Agilent 2100 Bioanalyzer, a standard tool for measuring RNA quality, which calculates the RNA Integrity Number (RIN). RNA integrity ranges from completely degraded RNA (RIN=1) to fully intact RNA (RIN=10). RIN=6 is a typical cutoff for downstream RNA sequencing. All extracted samples (n=23/23) had RIN > 6 (Fig. 4A), indicating sufficient stabilization of RNA with the homeRNAdrop. For general information on electropherograms generated by bioanalyzers and also electropherograms for samples in this study, see Figs. S8-S9.

**Figure 4.**
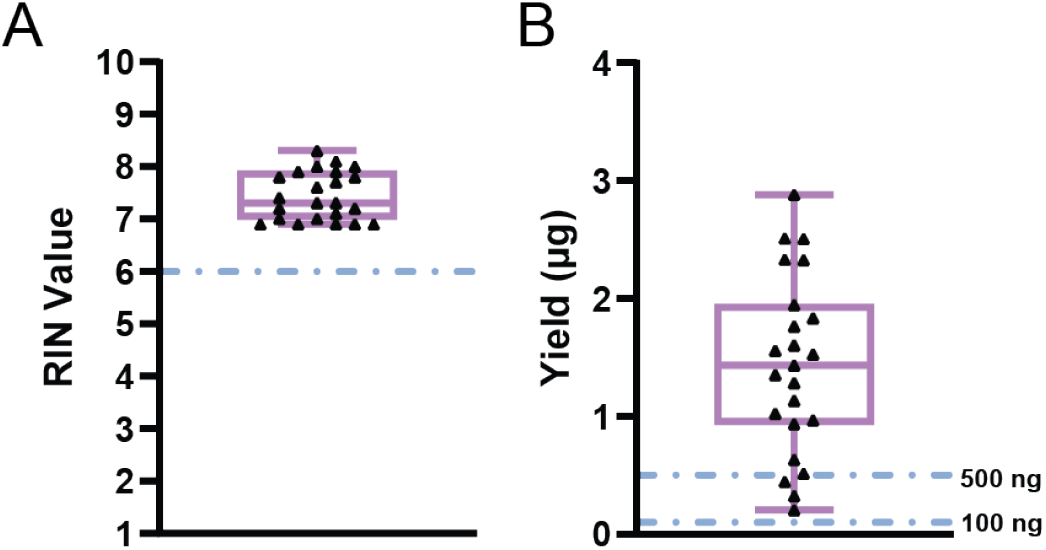
RNA integrity and yield of homeRNAdrop-stabilized blood samples from a nationwide remote pilot study. Both graphs are represented as standard box-and-whisker plots with each triangle representing a sample from a unique human participant (n=23 for both graphs). One sample was lost in transit by the courier (UPS), and one participant experienced failure to collect blood using the Tasso Mini device. (A) RNA integrity number (RIN) of homeRNAdrop samples obtained using an Agilent 2100 Bioanalyzer.

To quantify RNA yield, the RNA concentrations were measured on the Qubit Flex, a fluorometric assay specific to RNA. The concentrations were then converted to yield in micrograms and reported in Fig. 4B. Nearly all samples (n=20/23) had yields greater than 500 ng, a conservative estimate of RNA needed for RNA sequencing. Furthermore, all samples (n=23/23) had sufficient yields for targeted gene analysis (usually 100 ng or more recommended). For more information on RNA concentration, yield, and blood collection data for samples collected in pilot study, see Table S3 and Fig. S10.

Taken together, these results strongly indicate that homeRNAdrop is effective at stabilizing and preserving RNA in typical conditions for homeRNA-based studies. The dashed line represents a RIN of 6, a common cutoff for RNA sequencing technologies. (B) Yield measurements obtained using a Qubit Flex Fluorometer. The dashed lines represent 500 ng, a conservative cutoff for yield required for large-scale transcriptomic analysis, and 100 ng, a conservative cutoff for targeted gene analysis.

## Conclusion

homeRNAdrop is a new platform for remote RNA stabilization in blood, allowing for collection and RNA stabilization of up to 200 μL blood to occur in the same blood tube. It is designed to fit in the commercially available BD Microtainer tube, allowing homeRNAdrop to be compatible with multiple upper-arm blood collection devices, such as the Tasso Mini and the Tap Micro Select. Its cone-shaped channel allows for blood and stabilizer mixing, while also preventing the chemical stabilizer from spilling and coming into contact with a participant. Further, the homeRNAdrop makes the blood stabilization process more efficient than our other homeRNA platforms by eliminating the need for a separate stabilizer tube and can be easily modified to control the ratio of the stabilizer and blood sample, making it suitable for a host of applications.

Our pilot study of n=25 total participants shows that homeRNA drop is essentially painless (n=20/25) and easy to use (n=25 for the Tasso Mini and n=23/24 for the homeRNAdrop device), and sampling was quick (n=22/24 completed sampling in 15 minutes or less). The RNA extracted from these samples was of sufficient quality (mean RIN=7.4) and yield (mean yield=1.44 μg) for downstream gene expression analysis. Future work includes applying homeRNAdrop to longitudinal studies in laboratories that could benefit from a sample tube with a smaller storage footprint and simplified workflow compared to our previous homeRNA platforms. The homeRNAdrop insert design can also be modified for a variety of different use cases outside of RNA preservation, including analytes that might require a different ratio of blood:stabilizer.

## Acknowledgements

This publication was supported by the National Institutes of Health (NIH) through the National Institute of General Medical Sciences award number R35GM128648 (for in-lab developments, testing, and engineering), the National Center for Advancing Translational Sciences TL1TR002318 (ME, trainee), and the Schmidt Sciences Polymath Program (for the human subjects component of the research). The REDCap used for human subjects consent and enrollment is supported by the Institute of Translational Health Sciences, which is funded by the National Center for Advancing Translational Sciences of the National Institutes of Health under award number UL1TR002319. The content is solely the responsibility of the authors and does not necessarily represent the official views of the National Institutes of Health or other funding bodies. We would also like to thank the participants for their valued contributions.

## References

(1) Petrini, C.; Mannelli, C.; Riva, L.; Gainotti, S.; Gussoni, G. Decentralized Clinical Trials (DCTs): A Few Ethical Considerations. Front. Public Health 2022, 10, 1081150. 10.3389/fpubh.2022.1081150

(2) Martial, L. C.; Aarnoutse, R. E.; Schreuder, M. F.; Henriet, S. S.; Brüggemann, R. J. M.; Joore, M. A. Cost Evaluation of Dried Blood Spot Home Sampling as Compared to Conventional Sampling for Therapeutic Drug Monitoring in Children. PLOS ONE 2016, 11 (12), e0167433. 10.1371/journal.pone.0167433

(3) Koulman, A.; Prentice, P.; Wong, M. C. Y.; Matthews, L.; Bond, N. J.; Eiden, M.; Griffin, J. L.; Dunger, D. B. The Development and Validation of a Fast and Robust Dried Blood Spot Based Lipid Profiling Method to Study Infant Metabolism. Metabolomics 2014, 10 (5), 1018–1025. 10.1007/s11306-014-0628-z

(4) Trifonova, O. P.; Maslov, D. L.; Balashova, E. E.; Lokhov, P. G. Evaluation of Dried Blood Spot Sampling for Clinical Metabolomics: Effects of Different Papers and Sample Storage Stability. Metabolites 2019, 9 (11), 277. 10.3390/metabo9110277

(5) Fuller, G.; Njune Mouapi, K.; Joung, S.; Shufelt, C.; Van Den Broek, I.; Lopez, M.; Robinson, A.; Dhawan, S.; Mastali, M.; Spiegel, B.; Bairey Merz, N.; Van Eyk, J. E. Feasibility of Patient-Centric Remote Dried Blood Sampling: The Prediction, Risk, and Evaluation of Major Adverse Cardiac Events (PRE-MACE) Study. Biodemography Soc. Biol. 2020, 65 (4), 313–322. 10.1080/19485565.2020.1765735

(6) Li, K.; Naviaux, J. C.; Monk, J. M.; Wang, L.; Naviaux, R. K. Improved Dried Blood Spot-Based Metabolomics: A Targeted, Broad-Spectrum, Single-Injection Method. Metabolites 2020, 10 (3), 82. 10.3390/metabo10030082

(7) Lim, M. D. Dried Blood Spots for Global Health Diagnostics and Surveillance: Opportunities and Challenges. Am. J. Trop. Med. Hyg. 2018, 99 (2), 256–265. 10.4269/ajtmh.17-0889

(8) Baillargeon, K. R.; Mace, C. R. Microsampling Tools for Collecting, Processing, and Storing Blood at the Point-of-care. Bioeng. Transl. Med. 2023, 8 (2), e10476. 10.1002/btm2.10476

(9) Fredolini, C.; Dodig-Crnković, T.; Bendes, A.; Dahl, L.; Dale, M.; Albrecht, V.; Mattsson, C.; Thomas, C. E.; Torinsson Naluai Å.; Gisslen, M.; Beck, O.; Roxhed, N.; Schwenk, J. M. Proteome Profiling of Home-Sampled Dried Blood Spots Reveals Proteins of SARS-CoV-2 Infections. Commun. Med. 2024, 4 (1), 55. 10.1038/s43856-024-00480-4

(10) Wikström, F.; Olsson, C.; Palm, B.; Roxhed, N.; Backlund, L.; Schalling, M.; Beck, O. Determination of Lithium Concentration in Capillary Blood Using Volumetric Dried Blood Spots. J. Pharm. Biomed. Anal. 2023, 227, 115269. 10.1016/j.jpba.2023.115269

(11) Lenk, G.; Ullah, S.; Stemme, G.; Beck, O.; Roxhed, N. Evaluation of a Volumetric Dried Blood Spot Card Using a Gravimetric Method and a Bioanalytical Method with Capillary Blood from 44 Volunteers. Anal. Chem. 2019, 91 (9), 5558–5565. 10.1021/acs.analchem.8b02905

(12) Baillargeon, K. R.; Brooks, J. C.; Miljanic, P. R.; Mace, C. R. Patterned Dried Blood Spot Cards for the Improved Sampling of Whole Blood. ACS Meas. Sci. Au 2022, 2 (1), 31–38. 10.1021/acsmeasuresciau.1c00031

(13) Bybjerg-Grauholm, J.; Hagen, C. M.; Khoo, S. K.; Johannesen, M. L.; Hansen, C. S.; Bækvad-Hansen, M.; Christiansen, M.; Hougaard, D. M.; Hollegaard, M. V. RNA Sequencing of Archived Neonatal Dried Blood Spots. Mol. Genet. Metab. Rep. 2017, 10, 33–37. 10.1016/j.ymgmr.2016.12.004

(14) Bertoli-Avella, A. M.; Radefeldt, M.; Al-Ali, R.; Pardo, L. M.; Lemke, S.; Leubauer, A.; Polla, D. L.; Hörnicke, R.; Almeida, L. S.; Kandaswamy, K. K.; Beetz, C.; Pinto Basto, J.; Bauer, P. Beyond Genomics: Using RNA-Seq from Dried Blood Spots to Unlock the Clinical Relevance of Splicing Variation in a Diagnostic Setting. Eur. J. Hum. Genet. 2025, 33 (5), 614–623. 10.1038/s41431-025-01792-2

(15) Haack, A. J.; Lim, F. Y.; Kennedy, D. S.; Day, J. H.; Adams, K. N.; Lee, J. J.; Berthier, E.; Theberge, A. B. homeRNA: A Self-Sampling Kit for the Collection of Peripheral Blood and Stabilization of RNA. Anal. Chem. 2021, 93 (39), 13196–13203. 10.1021/acs.analchem.1c02008

(16) Eakman, M.; Stefanovic, F.; Berthier, J.; Robertson, I. H.; Knudsen, L. A.; Cardoso Carvalho, C.; Chen, K.; Atkeson, J.; Eichman, S.; Nguyen, S. H.; Craig, C. A.; Leong, K. M.; Chan, D. W. H.; Adams, K. N.; Olanrewaju, A. O.; Thongpang, S.; Nicholson, T. M.; Haberman, R. H.; Scher, J. U.; Berthier, E.; Theberge, A. B.; Haack, A. J. To the homeRNAmax: Developing an Improved Blood Self-Collection and Stabilization Platform for Remote Transcriptomic Studies. Anal. Chem. 2026, 98 (25), 18363–18372. 10.1021/acs.analchem.5c07772

(17) Lim, F. Y.; Kim, S.-Y.; Kulkarni, K. N.; Blazevic, R. L.; Kimball, L. E.; Lea, H. G.; Haack, A. J.; Gower, M. S.; Stevens-Ayers, T.; Starita, L. M.; Boeckh, M.; Hyrien, O.; Schiffer, J. T.; Theberge, A. B.; Waghmare, A. High-Frequency Home Self-Collection of Capillary Blood Correlates IFI27 Expression Kinetics with SARS-CoV-2 Viral Clearance. J. Clin. Invest. 2023, 133 (23), e173715. 10.1172/JCI173715

(18) Lim, F. Y.; Lea, H. G.; Dostie, A. M.; Kim, S.-Y.; van Neel, T. L.; Hassan, G. W.; Takezawa, M. G.; Starita, L. M.; Adams, K. N.; Boeckh, M.; Schiffer, J. T.; Hyrien, O.; Waghmare, A.; Berthier, E.; Theberge, A. B. homeRNA Self-Blood Collection Enables High-Frequency Temporal Profiling of Presymptomatic Host Immune Kinetics to Respiratory Viral Infection: A Prospective Cohort Study. EBioMedicine 2025, 112, 105531. 10.1016/j.ebiom.2024.105531

(19) Haack, A. J.; Brown, L. G.; Zeng, Y.; Khan, T.; Robertson, I. H.; Kennedy, D. S.; Adams, K. N.; MacDonald, J. W.; Bammler, T. K.; Stefanovic, F.; Moloney, K.; Stolarczuk, J. E.; Alizai, M. Y.; Hassan, G. W.; Lim, F. Y.; Chaussabel, D.; Walker, E. G.; Berthier, E.; Theberge, A. B. A Flexible and Responsive Remote Study Design to Assess Gene Expression Changes During Wildfire Smoke Exposure with homeRNA, an At-Home Blood Sampling Kit.

(20) Brown, L. G.; Wei, X.; Milton, L. A.; Alizai, M. Y.; MacDonald, J. W.; Bammler, T. K.; Zeng, Y.; Robertson, I.; Adams, K. N.; Toh, Y.-C.; Chaussabel, D.; Berthier, E.; Haack, A. J.; Theberge, A. B. From Home to Transcriptome: Comparing the Transcriptomic Profile of Induced Immune Response via Lipopolysaccharide Stimulation in homeRNA and Venous Blood. Anal. Chem. 2026, 98 (4), 2990–3001. 10.1021/acs.analchem.5c06040

(21) Brown, L. G.; Haack, A. J.; Kennedy, D. S.; Adams, K. N.; Stolarczuk, J. E.; Takezawa, M. G.; Berthier, E.; Thongpang, S.; Lim, F. Y.; Chaussabel, D.; Garand, M.; Theberge, A. B. At-Home Blood Collection and Stabilization in High Temperature Climates Using homeRNA. Front. Digit. Health 2022, 4, 903153. 10.3389/fdgth.2022.903153

(22) Stefanovic, F.; Brown, L. G.; MacDonald, J.; Bammler, T.; Rinchai, D.; Nguyen, S.; Zeng, Y.; Shinkawa, V.; Adams, K.; Chaussabel, D.; Berthier, E.; Haack, A. J.; Theberge, A. B. Your Blood Is Out for Delivery: Considerations of Shipping Time and Temperature on Degradation of RNA from Stabilized Whole Blood. Anal. Chem. 2025, 97 (3), 1635–1644. 10.1021/acs.analchem.4c04591

(23) Stefanovic, F.; Robertson, I.; Moloney, K.; Edelson, J.; Nguyen, S.; Shinkawa, V.; Uchimura, K.; Lin, A.; Le, L.; Tokihiro, J. C.; Takezawa, M. G.; Phan, D.; Schiffer, J. T.; Boeckh, M.; Adams, N.; Waghmare, A.; Errett, N. A.; Berthier, E.; Lim, F. Y.; Theberge, A. B. Research In Your Mailbox: Remote Blood Self-Sampling Enables Participation of Underserved Populations in Longitudinal Studies.

