## Supplementary Information 1 for "homeRNA Just Dropped! Streamlining Remote Blood Collection and RNA Stabilization for Smaller Sample Volumes with homeRNAdrop"

**Table of Contents:**


| | |
|---|---|
| **Figure S1:** Results of shipping tests | S1 |
| **Table S1:** Summary of results of shipping tests | S1 |
| **Figure S2:** Engineering drawings | S2 |
| **Figure S3:** Summary of evaporation test data | S3 |
| **Figure S4:** Instructions for Use | S4-S5 |
| **Figure S5:** Instructions for sample packaging and return shipment | S6 |
| **Figure S6:** Components of the homeRNAdrop kit | S7 |
| **Table S2:** Components of the homeRNAdrop kit | S7 |
| **Figure S7:** Stickers showing adequate sample stabilization | S8 |
| **Figure S8:** Bioanalyzer general profile and information | S8 |
| **Figure S9:** Bioanalyzer electropherograms | S19 |
| **Table S3:** Summary of yield and sample blood collection | S10-S11 |
| **Figure S10:** Blood levels of pilot study samples | S12 |
| **References** | S13 |

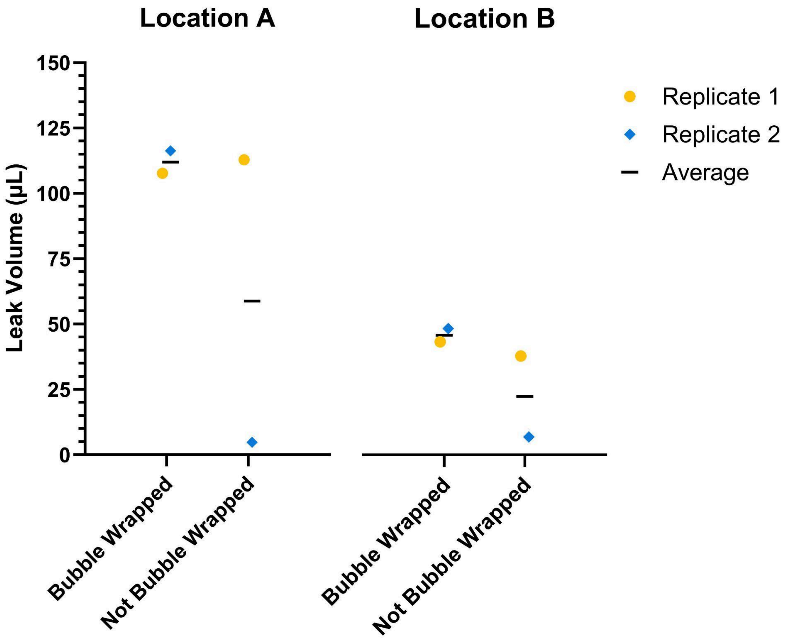


**Figure S1.** Results of preliminary shipping tests for homeRNAdrop device with and without applying bubble wrap to the outside of the cardboard box. Experiment was performed in duplicate; two complete kits were sent in separate packages to one lab member's house ("Location A") through UPS, and another two were sent to a different lab member's house ("Location B"), and shipped back to the lab. Both locations are in the Seattle area.

**Table S1.** Summary of fluid loss into top compartment from shipping tests described in Fig. S1.

| | Leak Volume (µL) | | | | | | |
|---|---|---|---|---|---|---|---|
| | Location A | | | Location B | | | |
| | Replicate 1 | Replicate 2 | Average | Replicate 1 | Replicate 2 | Average | Average (Across both locations) |
| Bubble Wrapped | 116.24 | 107.68 | 111.96 | 41.97 | 46.85 | 44.41 | 78.19 |
| Not Bubble Wrapped | 4.80 | 112.88 | 58.84 | 36.70 | 6.72 | 21.71 | 40.27 |

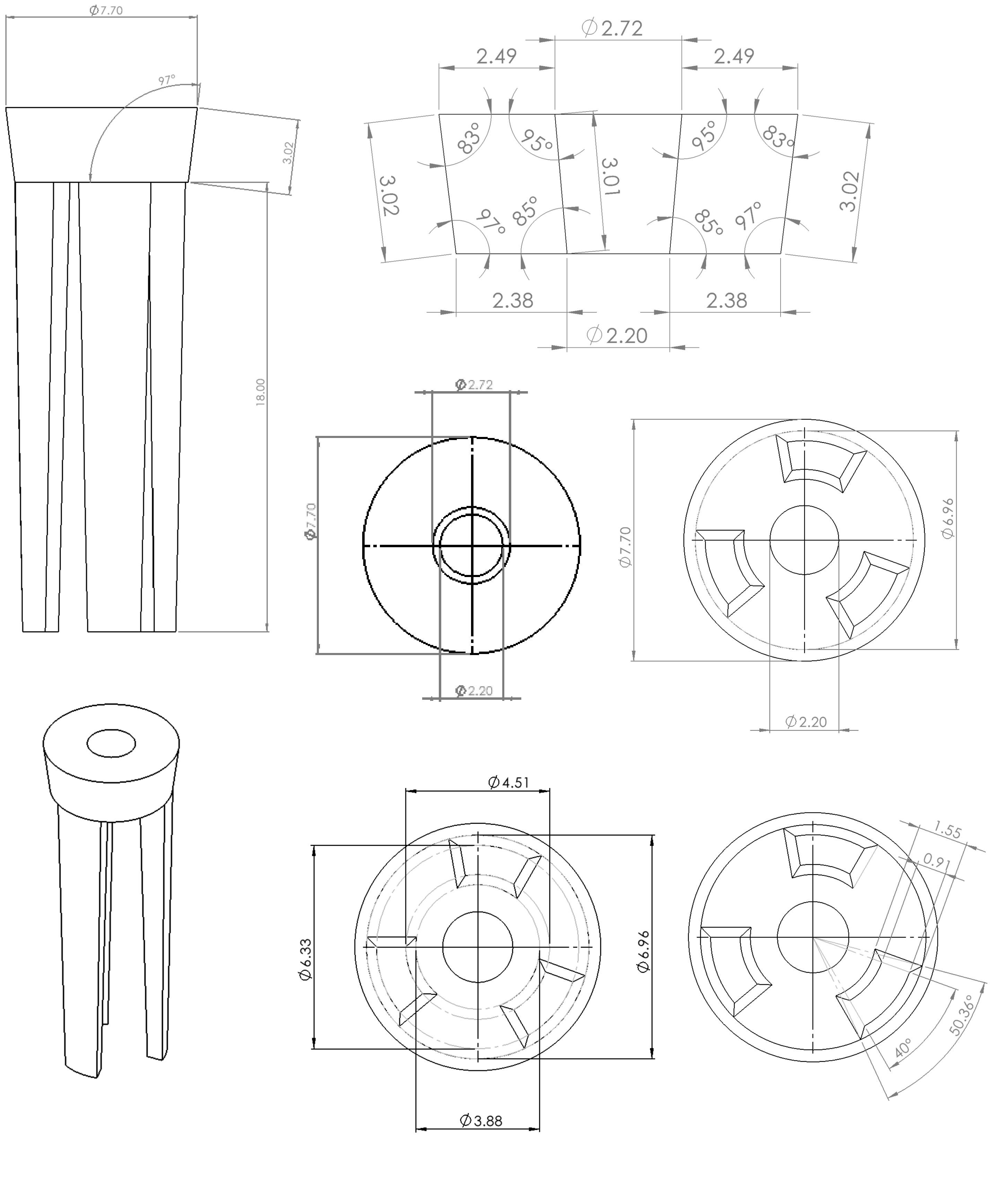


**Figure S2.** Engineering drawings for homeRNAdrop. All dimensions are in millimeters and legs were drafted 1 degree.

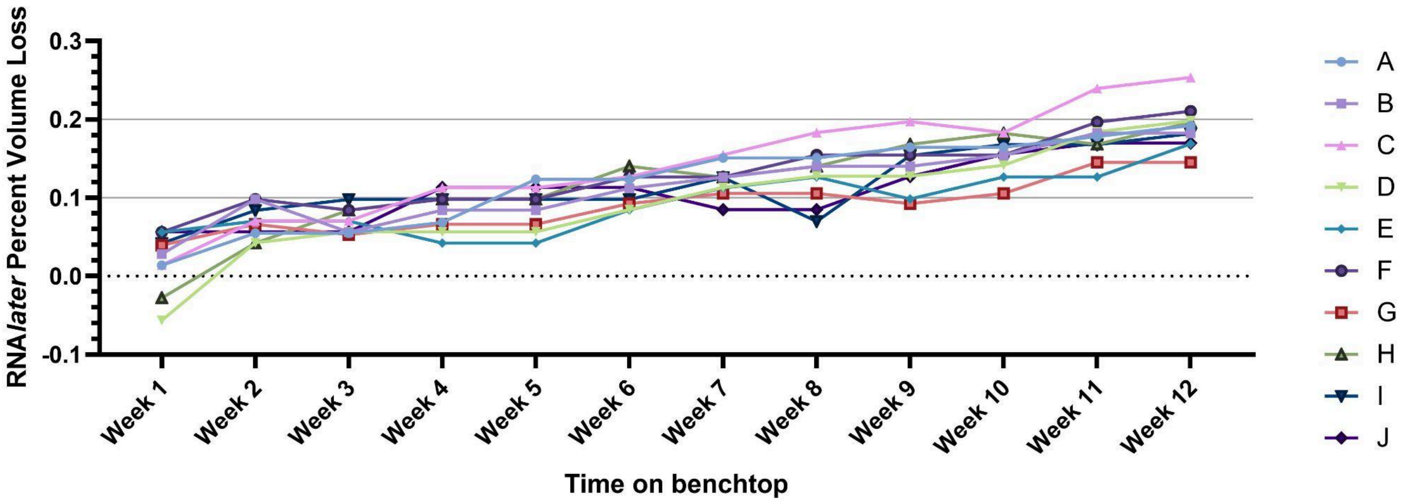


**Figure S3**. Graph of evaporation test data. Ten homeRNAdrop tubes were filled with 590 µL RNA*later* and stored upright at room temperature on a lab bench (~21-24 ℃). The mass was measured each week, and the percent loss was calculated. These data show minimal evaporation, with less than 0.3 % volume lost after 12 weeks for all samples. Replicates D and H reported negative loss after 1 week of room temperature storage; this was likely due to variance within the scale used to take measurements (the actual mass difference for D and H after one week of storage was < 0.0005 g).

### PREPARE FOR COLLECTION

**BEFORE YOU BEGIN**

Point your phone's camera here to watch an instructional video:

Or type this URL: tinyurl.com/3fvt7jwr

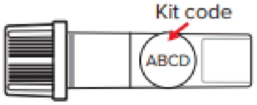


**1. Write down the sample code on the blood collection tube.** You will use this for step 16.

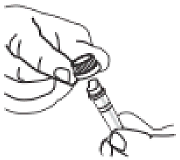

**2. Twist and remove cap from blood collection tube.** Set cap aside, you will use this in step 13.

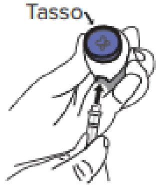


**3. Remove Tasso from box. Connect blood collection tube to Tasso device, and press until snug.**

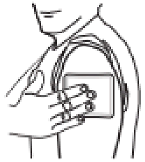

**4. Activate warmer by bending the silver disc. Knead to fully activate, and apply to upper arm for 2 minutes.**

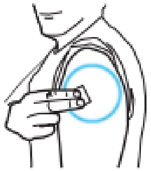

**5. Clean area with alcohol wipe and allow to dry.**

### COLLECT BLOOD USING TASSO DEVICE

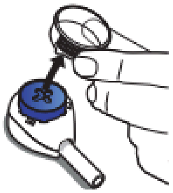

**6. Remove clear plastic cover over the blue button on the Tasso device.**

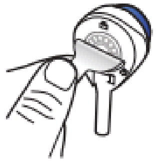

**7. Peel tab behind the Tasso device.** Keep the tube pointing down.

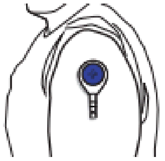

**8. Stick device to upper arm. Hang arm straight down at side.** Do not remove once it is on.

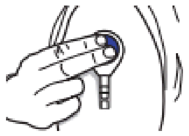

**9. Press button all the way down ONCE and release.** You may not feel anything.

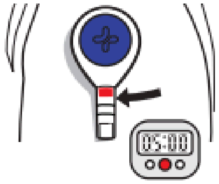


**10. Start a 5 minute timer.** Keep arm at your side. You may not see blood right away. It can take 1-2 minutes for blood to flow. Use a mirror or phone camera as needed to watch blood flow.

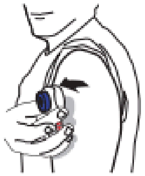

**11. After 5 minutes OR when the blood reaches the collar of the tube, *whichever comes first*, remove the device from your arm.**

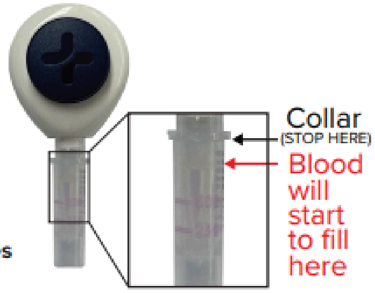


**NOTE: If you have not collected any blood skip to step 16.**

### STABILIZE AND PACKAGE

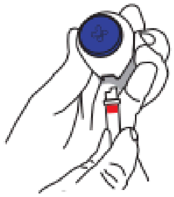

**12. Remove tube by twisting slightly and pulling down.** This may take a bit of finger strength.

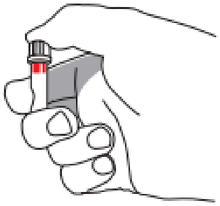

**13. Snap cap fully onto blood collection tube.**

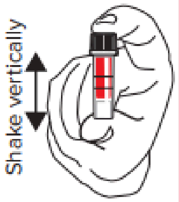


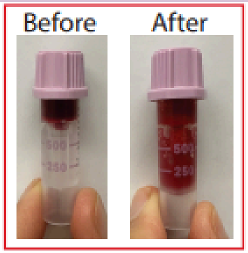


**14. Shake up and down *__vigorously__* until the fluid in the top and bottom compartments is the same color. For most participants, this takes about 10 seconds of *__vigorous__* shaking.**

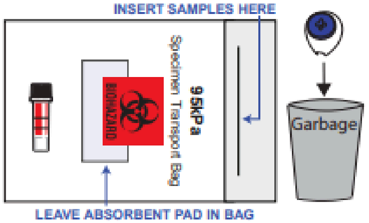


**15. Place blood sample in specimen bag, then put specimen bag inside the original box.** Remaining used materials, including the Tasso device and warmer, can be thrown away.

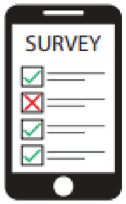


**16. Refer to email instructions for filling out the Online Survey and for mailing your sample back to the lab.**

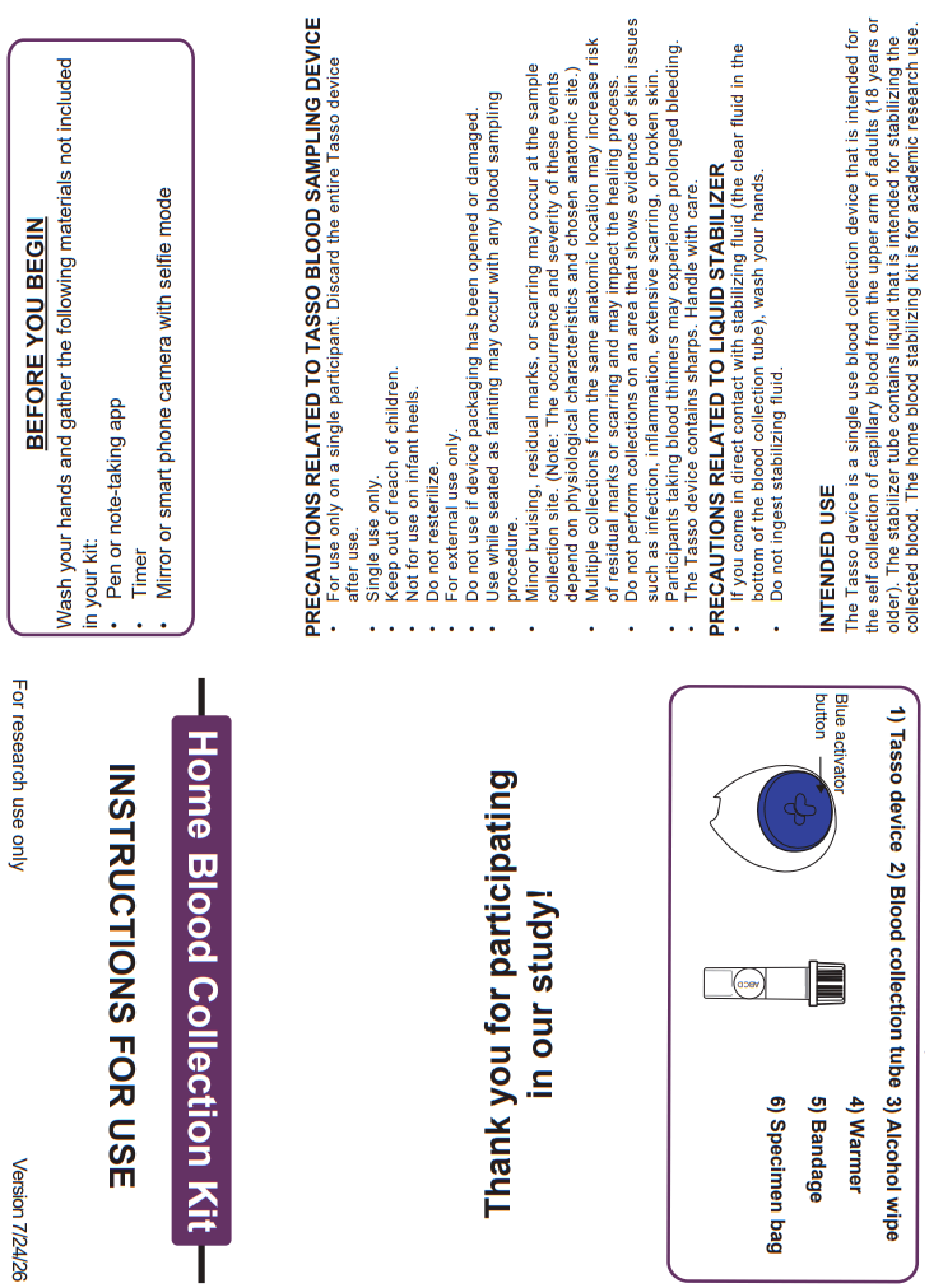

### BEFORE YOU BEGIN

Wash your hands and gather the following materials not included in your kit:

- Pen or note-taking app
- Timer
- Mirror or smart phone camera with selfie mode

**PRECAUTIONS RELATED TO TASSO BLOOD SAMPLING DEVICE**

- For use only on a single participant. Discard the entire Tasso device after use.
- Single use only.
- Keep out of reach of children.
- Not for use on infant heels.
- Do not resterilize.
- For external use only.
- Do not use if device packaging has been opened or damaged.
- Use while seated as fainting may occur with any blood sampling procedure.
- Minor bruising, residual marks, or scarring may occur at the sample collection site. (Note: The occurrence and severity of these events depend on physiological characteristics and chosen anatomic site.)
- Multiple collections from the same anatomic location may increase risk of residual marks or scarring and may impact the healing process.
- Do not perform collections on an area that shows evidence of skin issues such as infection, inflammation, extensive scarring, or broken skin.
- Participants taking blood thinners may experience prolonged bleeding.
- The Tasso device contains sharps. Handle with care.

**PRECAUTIONS RELATED TO LIQUID STABILIZER**

- If you come in direct contact with stabilizing fluid (the clear fluid in the bottom of the blood collection tube), wash your hands.
- Do not ingest stabilizing fluid.

**INTENDED USE**

The Tasso device is a single use blood collection device that is intended for the self collection of capillary blood from the upper arm of adults (18 years or older). The stabilizer tube contains liquid that is intended for stabilizing the collected blood. The home blood stabilizing kit is for academic research use.

**STORAGE**

Store at 15 - 30ºC (60 - 80ºF) in a dry place.

**Home Blood Collection Kit**

**INSTRUCTIONS FOR USE**

**Thank you for participating in our study!**

For research use only

Version 7/24/26

**KIT CONTENTS**

Make sure your kit contains all components listed below

**1) Tasso device 2) Blood collection tube 3) Alcohol wipe**

**4) Warmer**

**5) Bandage**

**6) Specimen bag**



**Figure S4.** homeRNAdrop instructions for use (IFU). Adobe editable .pdf is included as an attachment in the Supplemental Information.

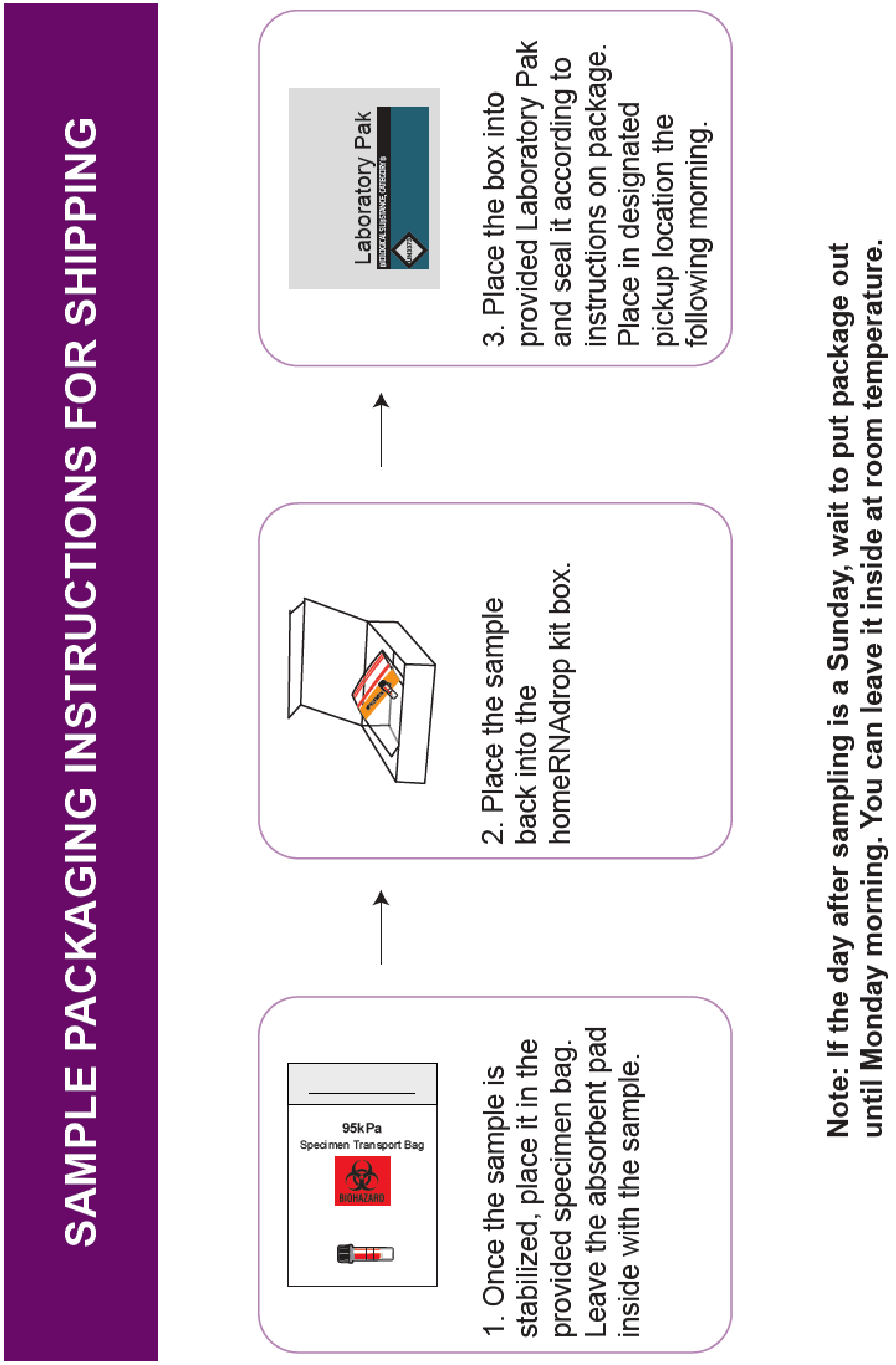


**Figure S5.** Instructions for sample packaging and return shipment. Adobe editable .pdf is included as an attachment in the Supplemental Information.

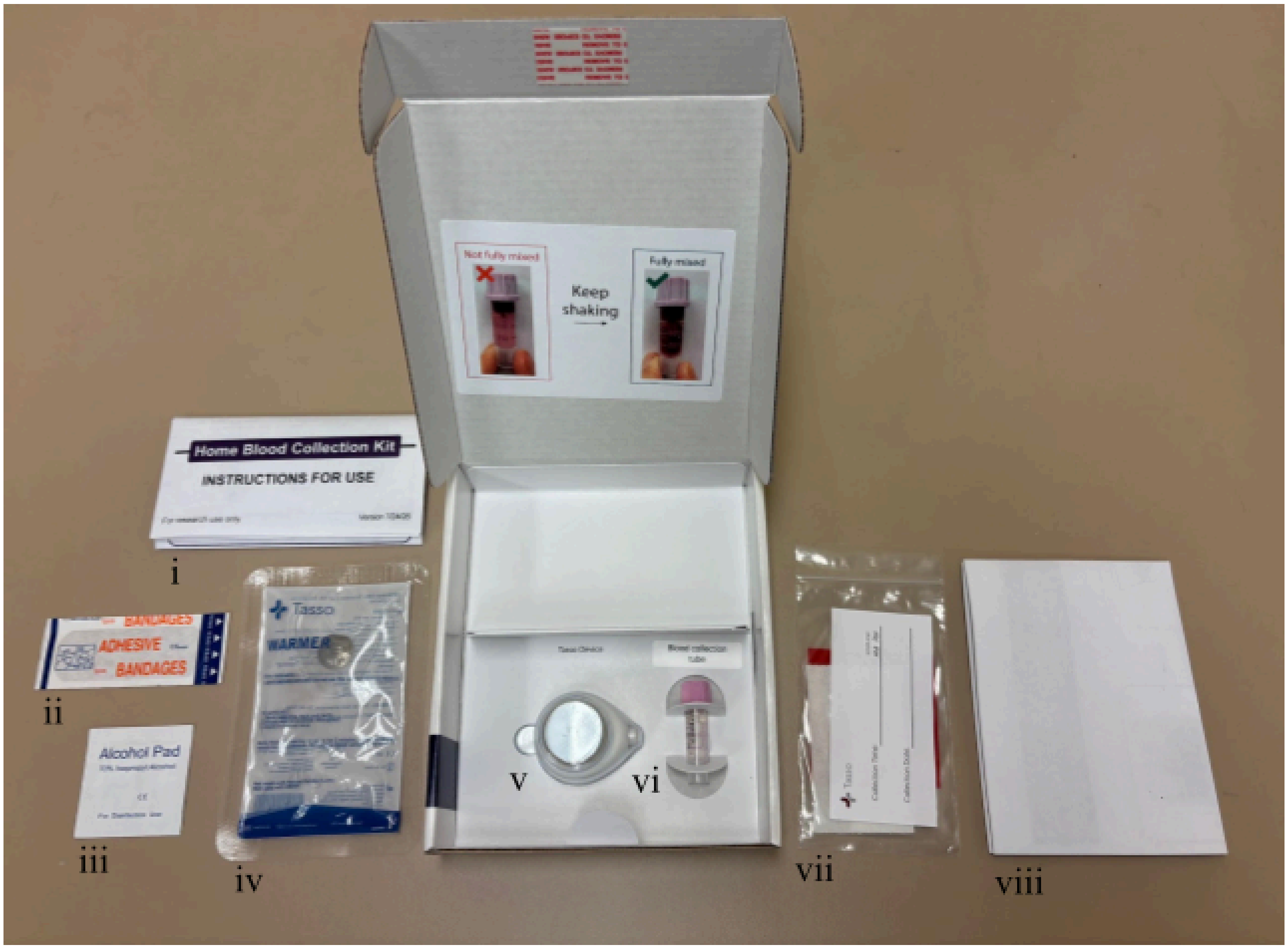


**Figure S6.** Components of the homeRNAdrop kit. Kit components include i) instructions for use, ii) sterile bandage, iii) alcohol wipe, iv) warmer for warming the arm prior to application of the Tasso Mini device, v) Tasso Mini device, vi) BD Microtainer blood tube fitted with homeRNAdrop insert and filled with 590 µL RNAlater, vii) sample return bag, viii) shipping return instructions.

**Table S2. Components of the homeRNAdrop kit.**

| Kit component | Manufacturer(s) | Quantity |
|---|---|---|
| Sterile Tasso Mini blood collection device | Tasso, Inc. | 1 |
| homeRNAdrop device | Our lab | 1 |
| BD Microtainer® blood tube (K2EDTA coated) | BD | 1 |
| Instant heat pack (product number ACC0014) | Tasso, Inc. | 1 |
| Sterile alcohol wipe | Covidien | 1 |
| Sterile bandage | McKesson | 1 |
| Specimen transport bag with absorbent pad | Path-Tec | 1 |
| Shipping return instructions | Our lab (see page S5 of SI) | 1 |
| Instructions for use | Our lab (see page S3-S4 of SI) | 1 |
| Blood-stabilizer mixing instruction sticker | Our lab (see page S7 of SI) | 1 |

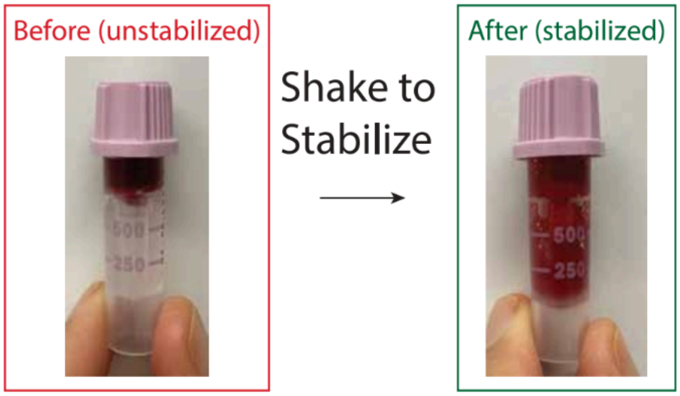


**Figure S7.** Stickers showing adequately stabilized blood samples. These stickers were placed on the inside of the box lid as shown in Figure S3 above. Labels used are Avery 5168. Adobe editable .pdf is included as an attachment in the Supplemental Information.

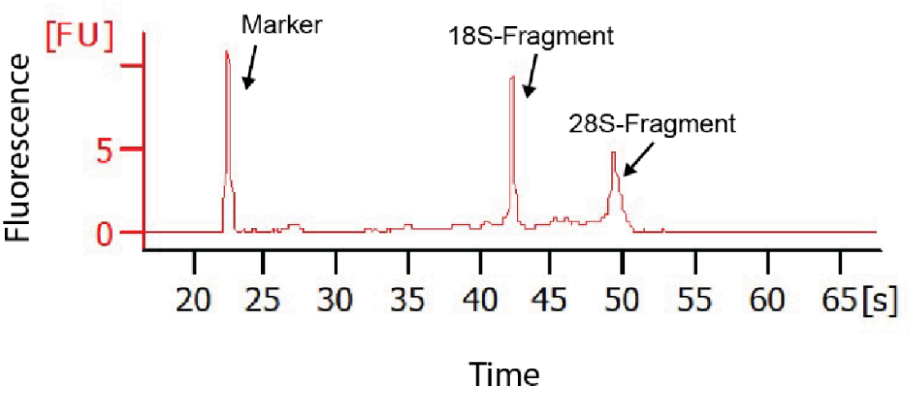


**Figure S8.** General information on Bioanalyzer profile. Electrophoretogram annotated with the marker and two other major peaks that help determine RIN including the 18-S fragment peak, and the 28-S fragment peak. More information on interpreting bioanalyzer data, including examples of electrophoretograms obtained from samples with various RIN values can be found in Schroeder 2006[1]. Figure and caption reproduced from Haack, Lim 2021[2] Supporting Information with permission.

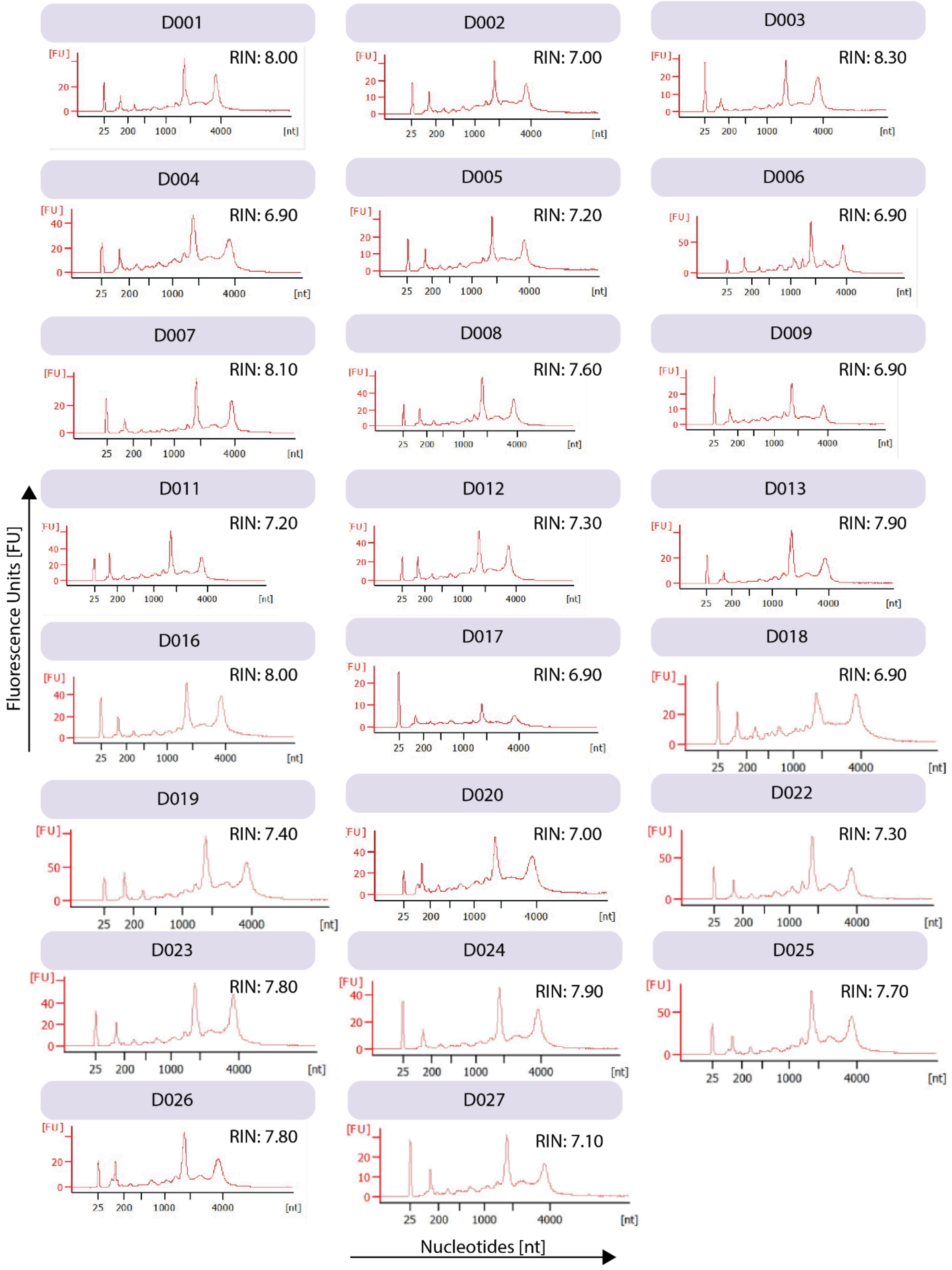


**Figure S9**. Agilent 2100 Bioanalyzer electropherograms for isolated RNA from homeRNAdrop-stabilized samples. All samples were first diluted 1:20 and analyzed with the Pico kit. Samples D003, D004, D013, D020, and D026 were rerun in duplicate using the Pico kit at 1:5 dilution due to low concentration. If RIN was different between the two replicates, the lowest RIN was reported here and in the manuscript.

**Table S3**. homeRNAdrop RNA concentration, yield, and data on blood collection. The concentrations were obtained on the Qubit Flex and are multiplied by 50 µL (elution volume). For the final yields reported in the manuscript, the yields from elution 1 and elution 2 are added together. Concentrations that were too low to be within the quantitative range of the Qubit High Sensitivity RNA assay were reported and used in final yield calculations as 0.00 ng/µL.

| Sample ID | Elution number | Concentration (ng/µL) | Yield (ng) | Time on arm (min.) | Blood level | Approximate blood volume (µL) |
|---|---|---|---|---|---|---|
| D001 | 1 | 22.7 | 1135 | 3-5 | Level B | 200 |
| | 2 | 7.81 | 391 | | | |
| D002 | 1 | 24.4 | 1220 | 3-5 | Above level B | > 200 |
| | 2 | 4.26 | 213 | | | |
| D003 | 1 | 6.52 | 326 | 3-5 | Level B | 200 |
| | 2 | 0.00 | 0 | | | |
| D004 | 1 | 15.7 | 785 | 3-5 | Level B | 200 |
| | 2 | 3.03 | 152 | | | |
| D005 | 1 | 43.6 | 2180 | < 3 | Level B | 200 |
| | 2 | 2.95 | 148 | | | |
| D006 | 1 | 55.4 | 2770 | 3-5 | Level A | 100 |
| | 2 | 2.18 | 109 | | | |
| D007 | 1 | 18.4 | 920 | < 3 | Level B | 200 |
| | 2 | 2.05 | 103 | | | |
| D008 | 1 | 33.0 | 1650 | < 3 | Level B | 200 |
| | 2 | 3.64 | 182 | | | |
| D009 | 1 | 12.6 | 630 | > 5 | Level A | 100 |
| | 2 | 0.00 | 0 | | | |
| D011 | 1 | 4.09 | 205 | 3-5 | Level B | 200 |
| | 2 | 0.00 | 0 | | | |
| D012 | 1 | 30.6 | 1530 | 3-5 | Level B | 200 |

| | | | | | | |
|---|---|---|---|---|---|---|
| | 2 | 4.64 | 232 | | | |
| D013 | 1 | 8.91 | 446 | 3-5 | Above level B | > 200 |
| | 2 | 0.00 | 0 | | | |
| D016 | 1 | 28.7 | 1435 | 3-5 | Level A | 100 |
| | 2 | 2.43 | 122 | | | |
| D017 | 1 | 15.9 | 795 | 3-5 | Above level B | > 200 |
| | 2 | 6.71 | 336 | | | |
| D018 | 1 | 37.9 | 1895 | < 3 | Level A | 100 |
| | 2 | 12.3 | 615 | | | |
| D019 | 1 | 47.7 | 2385 | < 3 | Above level B | > 200 |
| | 2 | 2.37 | 119 | | | |
| D020 | 1 | 17.1 | 855 | 3-5 | Level B | 200 |
| | 2 | 2.21 | 111 | | | |
| D022 | 1 | 29.6 | 1480 | 3-5 | Level B | 200 |
| | 2 | 2.43 | 122 | | | |
| D023 | 1 | 36.8 | 1840 | 3-5 | Level B | 200 |
| | 2 | 2.10 | 105 | | | |
| D024 | 1 | 24.1 | 1205 | 3-5 | Level B | 200 |
| | 2 | 2.93 | 147 | | | |
| D025 | 1 | 41.0 | 2050 | < 3 | Level A | 100 |
| | 2 | 5.53 | 277 | | | |
| D026 | 1 | 10.3 | 515 | 3-5 | Level B | 200 |
| | 2 | 0.00 | 0 | | | |
| D027 | 1 | 22.5 | 1125 | < 3 | Above level B | >200 |
| | 2 | 3.11 | 156 | | | |

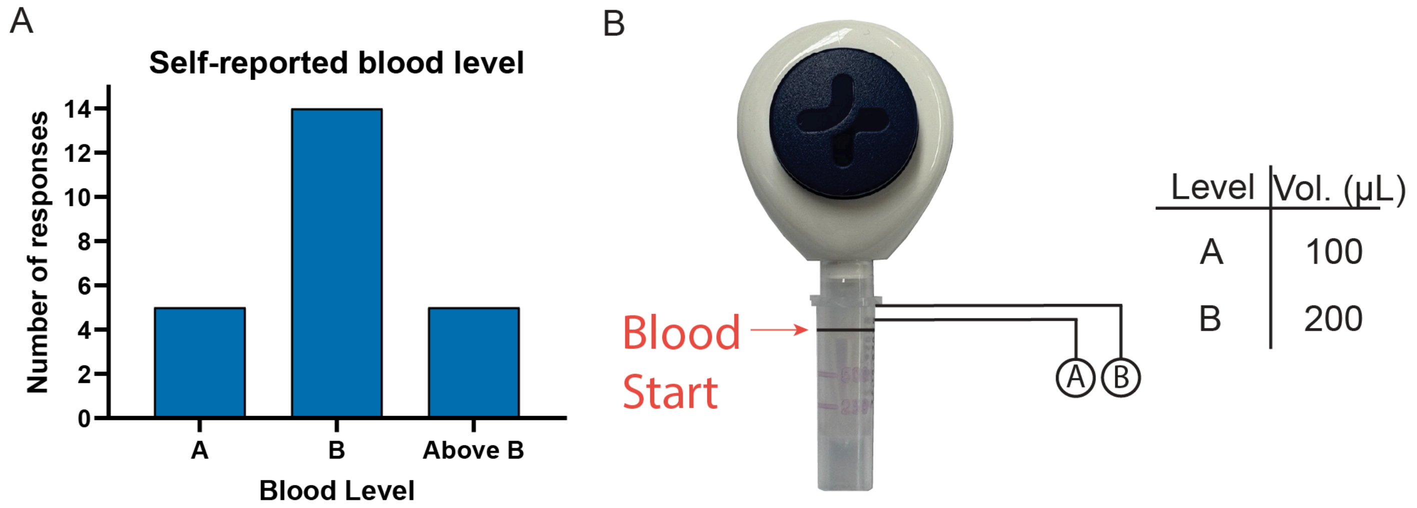


**Figure S10.** Participant-reported blood levels of samples collected using homeRNAdrop for the nationwide remote pilot study. (A) Bar graph showing distribution of blood levels reported by participants of the pilot study reported in this manuscript. (B) Image of Tasso Mini with homeRNAdrop device with annotated levels, and table showing the volumes represented by the levels. The image of the Tasso Mini is included in participant surveys as guidance for reporting collected blood level.

**References:**


(1) Schroeder, A.; Mueller, O.; Stocker, S.; Salowsky, R.; Leiber, M.; Gassmann, M.; Lightfoot, S.; Menzel, W.; Granzow, M.; Ragg, T. The RIN: An RNA Integrity Number for Assigning Integrity Values to RNA Measurements. *BMC Mol. Biol.* **2006**, *7* (1), 3. https://doi.org/10.1186/1471-2199-7-3

(2) Haack, A. J.; Lim, F. Y.; Kennedy, D. S.; Day, J. H.; Adams, K. N.; Lee, J. J.; Berthier, E.; Theberge, A. B. homeRNA: A Self-Sampling Kit for the Collection of Peripheral Blood and Stabilization of RNA. *Anal. Chem.* **2021**, *93* (39), 13196–13203. https://doi.org/10.1021/acs.analchem.1c02008
