## Supplementary Information 2 for "homeRNA Just Dropped! Streamlining Remote Blood Collection and RNA Stabilization for Smaller Sample Volumes with homeRNAdrop": homeRNAdrop_inside box sticker.pdf

Not fully mixed

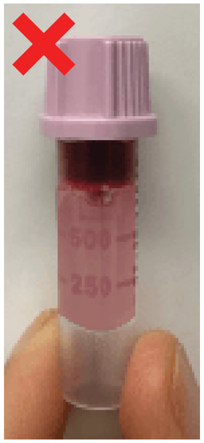

Keep shaking →

Fully mixed

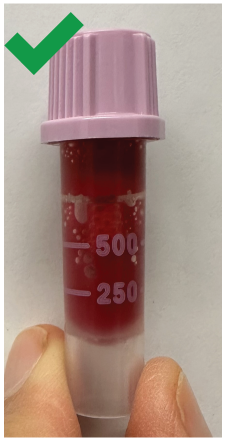


Not fully mixed

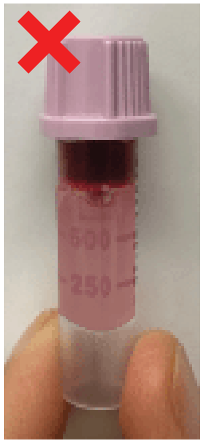

Keep shaking →

Fully mixed

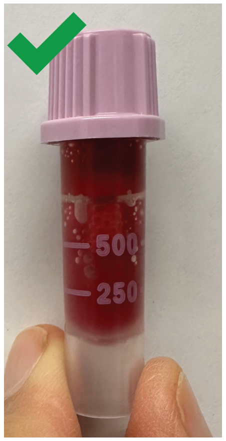


Not fully mixed

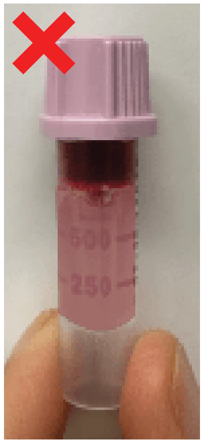

Keep shaking →

Fully mixed

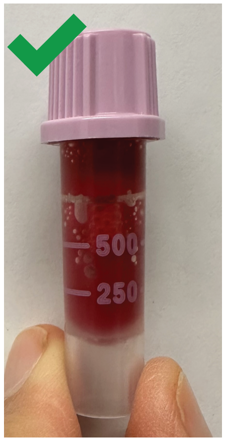


Not fully mixed

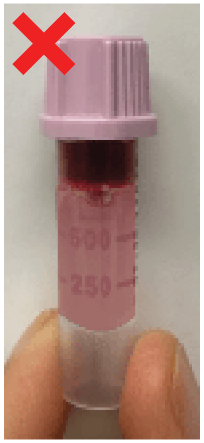

Keep shaking →

Fully mixed

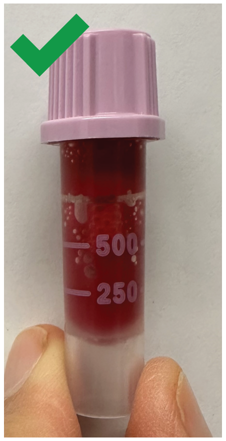
