## Supplementary Information 2 for "homeRNA Just Dropped! Streamlining Remote Blood Collection and RNA Stabilization for Smaller Sample Volumes with homeRNAdrop": homeRNAdrop_instructional email.pdf

Hi [prefname],

We have shipped your kit to you! You should receive the kit through UPS in the next day or two, but you will receive delivery updates from UPS through your email.

Please complete the following steps within 1 - 2 days after receiving your kit:

### 1. Collect the Sample

- Please watch the provided video before you begin sampling
  - Video is linked to QR code in physical instructions, and can be accessed through this link: https://youtu.be/T5U8ALbxSTE
- Follow printed instructions to collect your sample

### 2. Complete Survey

Next, please complete this survey after you are done with the sample collection:

[survey-link]

If the link above does not work, try copying the link below into your web browser:
[survey-url]

This link is unique to you and should not be forwarded to others.

### 3. Prepare Sample for Shipping

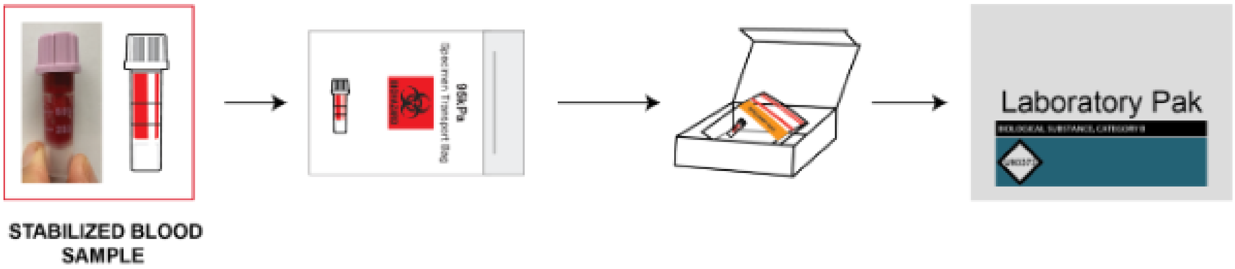


- Once the sample is packaged in the kit box, place it inside the pre-labeled UPS LabPak

    - Seal LabPak by removing the adhesive strip and pressing the flaps together
    - **Note:** Your kit will be shipped to you wrapped in bubble wrap; you do **<u>not</u>** need to wrap your kit to return your sample to the lab. The bubble wrap can be discarded.
- Keep sample indoors overnight at room temperature
- Place LabPak at the designated pickup location the following morning by 9:00am
    - **Note:** If the next day is Sunday, please leave package inside until Monday morning

There is nothing you need to do to schedule the pickup; we automatically schedule them after you complete the survey. You will receive an additional email letting you know what day the pickup is scheduled for.

Thank you again for your participation and please let us know if you have questions or need assistance!

All the best,

BCME Study Team

University of Washington
