## Supplementary Information 2 for "homeRNA Just Dropped! Streamlining Remote Blood Collection and RNA Stabilization for Smaller Sample Volumes with homeRNAdrop": homeRNAdrop_Instructions for Use.pdf

#### PREPARE FOR COLLECTION

##### BEFORE YOU BEGIN

Point your phone's camera here to watch an instructional video:

Or type this URL: tinyurl.com/3fvt7jwr

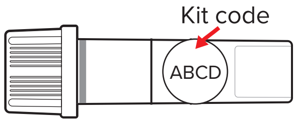


**1. Write down the sample code on the blood collection tube.** You will use this for step 16.

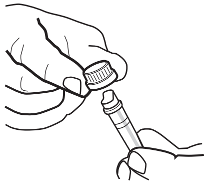

**2. Twist and remove cap from blood collection tube.** Set cap aside, you will use this in step 13.

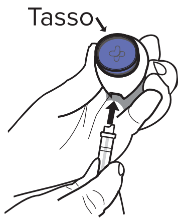


**3. Remove Tasso from box. Connect blood collection tube to Tasso device, and press until snug.**

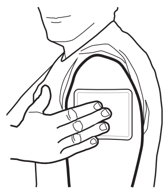

**4. Activate warmer by bending the silver disc. Knead to fully activate, and apply to upper arm for 2 minutes.**

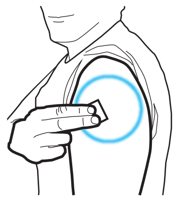

**5. Clean area with alcohol wipe and allow to dry.**

#### COLLECT BLOOD USING TASSO DEVICE

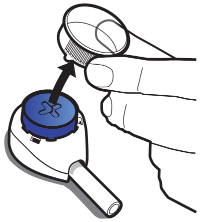

**6. Remove clear plastic cover over the blue button on the Tasso device.**

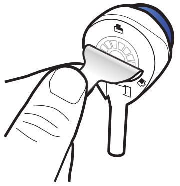

**7. Peel tab behind the Tasso device.** Keep the tube pointing down.

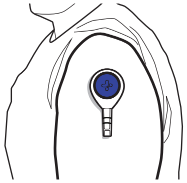

**8. Stick device to upper arm. Hang arm straight down at side.** Do not remove once it is on.

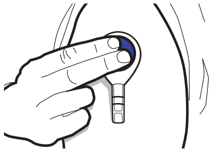

**9. Press button all the way down ONCE and release.** You may not feel anything.

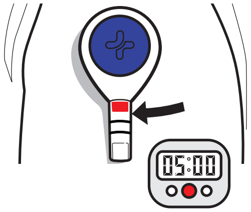


**10. Start a 5 minute timer.** Keep arm at your side. You may not see blood right away. It can take 1-2 minutes for blood to flow. Use a mirror or phone camera as needed to watch blood flow.

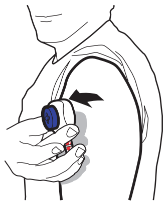

**11. After 5 minutes OR when the blood reaches the collar of the tube, *whichever comes first*, remove the device from your arm.**

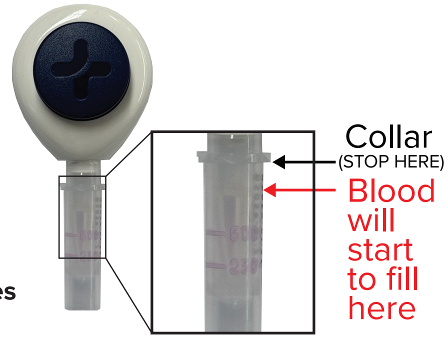


**NOTE: If you have not collected any blood skip to step 16.**

#### STABILIZE AND PACKAGE

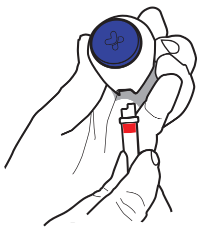

**12. Remove tube by twisting slightly and pulling down.** This may take a bit of finger strength.

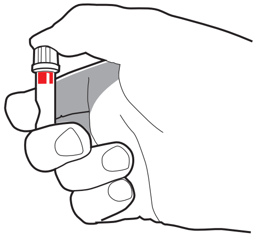

**13. Snap cap fully onto blood collection tube.**

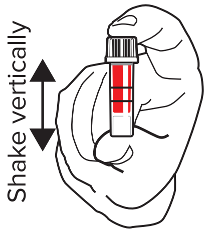


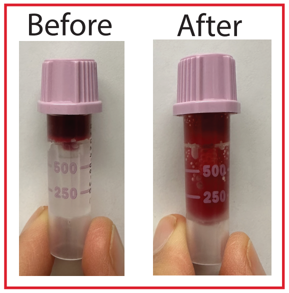


**14. Shake up and down *vigorously* until the fluid in the top and bottom compartments is the same color. For most participants, this takes about 10 seconds of *vigorous* shaking.**

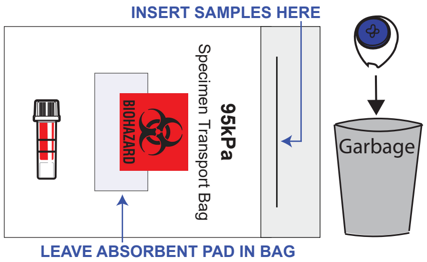


**15. Place blood sample in specimen bag, then put specimen bag inside the original box.** Remaining used materials, including the Tasso device and warmer, can be thrown away.

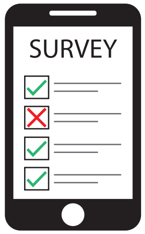


**16. Refer to email instructions for filling out the Online Survey and for mailing your sample back to the lab.**

### Home Blood Collection Kit

#### INSTRUCTIONS FOR USE

For research use only

Version 7/17/26

#### KIT CONTENTS

Make sure your kit contains all components listed below

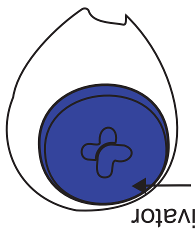


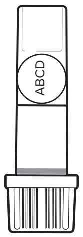


1) Tasso device 2) Blood collection tube 3) Alcohol wipe
4) Warmer
5) Bandage
6) Specimen bag

**Thank you for participating in our study!**

#### BEFORE YOU BEGIN

Wash your hands and gather the following materials not included in your kit:

- Pen or note-taking app
- Timer
- Mirror or smart phone camera with selfie mode

#### PRECAUTIONS RELATED TO TASSO BLOOD SAMPLING DEVICE

- For use only on a single participant. Discard the entire Tasso device after use.
- Single use only.
- Keep out of reach of children.
- Not for use on infant heels.
- Do not resterilize.
- For external use only.
- Do not use if device packaging has been opened or damaged.
- Use while seated as fainting may occur with any blood sampling procedure.
- Minor bruising, residual marks, or scarring may occur at the sample collection site. (Note: The occurrence and severity of these events depend on physiological characteristics and chosen anatomic site.)
- Multiple collections from the same anatomic location may increase risk of residual marks or scarring and may impact the healing process.
- Do not perform collections on an area that shows evidence of skin issues such as infection, inflammation, extensive scarring, or broken skin.
- Participants taking blood thinners may experience prolonged bleeding.
- The Tasso device contains sharps. Handle with care.

#### PRECAUTIONS RELATED TO LIQUID STABILIZER

- If you come in direct contact with stabilizing fluid (the clear fluid in the bottom of the blood collection tube), wash your hands.
- Do not ingest stabilizing fluid.

#### INTENDED USE

The Tasso device is a single use blood collection device that is intended for the self collection of capillary blood from the upper arm of adults (18 years or older). The stabilizer tube contains liquid that is intended for stabilizing the collected blood. The home blood stabilizing kit is for academic research use.

#### STORAGE

Store at 15 - 30ºC (60 - 80ºF) in a dry place.
