## Supplementary Information 2 for "homeRNA Just Dropped! Streamlining Remote Blood Collection and RNA Stabilization for Smaller Sample Volumes with homeRNAdrop": homeRNAdrop_shipping_instructions.pdf

### SAMPLE PACKAGING INSTRUCTIONS FOR SHIPPING

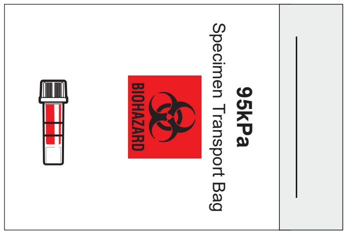


1. Once the sample is stabilized, place it in the provided specimen bag. Leave the absorbent pad inside with the sample.

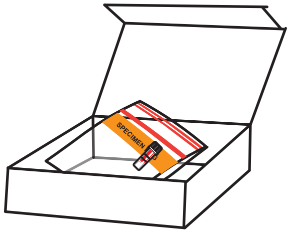

2. Place the sample back into the homeRNAdrop kit box.

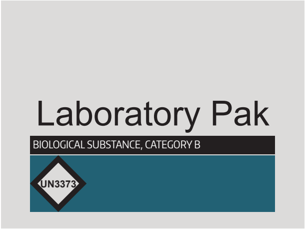


3. Place the box into provided Laboratory Pak and seal it according to instructions on package. Place in designated pickup location the following morning.

**Note: If the day after sampling is a Sunday, wait to put package out until Monday morning. You can leave it inside at room temperature.**
