## Supplementary Information 2 for "homeRNA Just Dropped! Streamlining Remote Blood Collection and RNA Stabilization for Smaller Sample Volumes with homeRNAdrop": homeRNAdrop_usability survey.pdf

### homeRNAdrop Kit Survey

Please complete the survey below.

Thank you!

---

What is the kit code?

Note: the kit code will consist of letters and numbers, and will look like D123.

__________________________________

---

Kit code as corrected by Study Team member (if needed):

__________________________________

---

Did you experience any issues with the Tasso blood collection device?

- ◯ Yes
- ◯ No

---

Please describe any issues you had with the Tasso blood collection device:

__________________________________________

---

Did you experience any pain when using the Tasso device today?

- ◯ No pain
- ◯ Mild pain
- ◯ Moderate pain
- ◯ Severe pain
- ◯ Very severe pain

---

Were you able to collect any blood?

- ◯ Yes
- ◯ No

---

If you have any comments or suggestions for the kit instructions, please use this space:

__________________________________________

---

Approximately how long (in minutes) did it take you to use the homeRNAdrop kit today?

- ◯ Less than 5 minutes
- ◯ 5-10 minutes
- ◯ 11-15 minutes
- ◯ 16-20 minutes
- ◯ More than 20 minutes

---

While using the homeRNAdrop kit today, approximately how long (in minutes) did you leave the Tasso blood collection device on your arm?

- ◯ Less than 1 minute
- ◯ Less than 3 minutes
- ◯ 3-5 minutes
- ◯ More than 5 minutes
- ◯ I am unsure

---

Based on the image below, how much blood did you roughly collect today? Choose the closest Level.

- ◯ Level A
- ◯ Level B
- ◯ Above level B (note: blood level B is the target stop point)

(Please just note as accurately as possible how much blood you collected, regardless of the level.)

---

Did you experience any issues mixing your blood with the clear fluid in the tube by shaking?

- ◯ Yes
- ◯ No

---

Please describe any issues you had:

__________________________________________

**How easy was it to use the components of the kit listed below?**

| | Very easy | Somewhat easy | Neither easy nor difficult | Somewhat difficult | Very difficult |
|---|---|---|---|---|---|
| Kit Instructions | ○ | ○ | ○ | ○ | ○ |
| Tasso Device | ○ | ○ | ○ | ○ | ○ |
| Blood Mixing | ○ | ○ | ○ | ○ | ○ |

Do you have any other thoughts or comments regarding your experience with this kit? (optional) ________________________________________
